# DNAmod: the DNA modification database

**DOI:** 10.1101/071712

**Authors:** Ankur Jai Sood, Coby Viner, Michael M. Hoffman

**Author notes:** Ankur Jai Sood and Coby Viner contributed equally to this work.

## Abstract

Covalent DNA modifications, such as 5-methylcytosine (5mC), are increasingly the focus of numerous research programs. In eukaryotes, both 5mC and 5-hydroxymethylcytosine (5hmC) are now recognized as stable epigenetic marks, with diverse functions. Bacteria, archaea, and viruses contain various other modified DNA nucleobases. Numerous databases describe RNA and histone modifications, but no database specifically catalogues DNA modifications, despite their broad importance in epigenetic regulation. To address this need, we have developed DNAmod: the DNA modification database.

DNAmod is an open-source database (https://dnamod.hoffmanlab.org) that catalogues DNA modifications and provides a single source to learn about their properties. DNAmod provides a web interface to easily browse and search through these modifications. The database annotates the chemical properties and structures of all curated modified DNA bases, and a much larger list of candidate chemical entities. DNAmod includes manual annotations of available sequencing methods, descriptions of their occurrence in nature, and provides existing and suggested nomenclature. DNAmod enables researchers to rapidly review previous work, select mapping techniques, and track recent developments concerning modified bases of interest.

## Introduction

A rapidly growing body of research is continuing to reveal numerous gene-regulatory effects of covalent DNA modifications, such as 5-methylcytosine (5mC). We now recognize 5mC as a stable epigenetic mark and as having diverse functions beyond transcriptional repression^11^. An increasing number of studies demonstrate the importance of other cytosine modifications, such as 5-hydroxymethyl-cytosine (5hmC), 5-formylcytosine (5fC), and 5-carboxylcytosine (5caC)^2,8,25,41,44^. More recently, three analogous modifications of thymine were found to occur in mammals^36,51^ and can now largely be sequenced^18^. *N*^6^-methyladenine, previously thought to mainly occur as an RNA modification in eukaryotes, has now been found in the DNA of multiple eukaryotes^23^. Bacteria, archaea, and especially bacteriophages have long been known to harbor a diverse array of modified bases^17,49^. Their genomes can also have hypermodified bases—modified DNA bases that substitute for the unmodified base in many positions genome-wide^16,49^.

Multiple databases profile RNA modifications^3,7,52^ and human histone modifications^54^, but no database catalogues DNA modifications systematically. Some databases include particular classes of DNA modifications^42^. These include restriction endonucleases and DNA methyltransferases in REBASE^39^; methylation databases, like MethDB^1^; databases including DNA metabolic pathways, such as KEGG^26^; and those focused on DNA damage and repair, like REPAIRtoire^29^.

Since DNA modifications are a key aspect of epigenetic regulation, there is a pressing need to organize them in a single location. We have accordingly created DNAmod: the DNA modification database (https://dnamod.hoffmanlab.org). DNAmod is the first database to comprehensively catalogue DNA modifications and provides a single resource to launch an investigation of their properties.

## Database construction and visualization

DNAmod consists of two components: a relational database back-end and a web interface frontend. We used the Chemical Entities of Biological Interest (ChEBI) database^12,21^ to seed the DNAmod database. We imported a nucleobase-related subset of ChEBI, consisting of chemical entities and related annotations. We performed queries against the entities to construct a set of candidate DNA modifications for DNAmod, retaining most of these as a separate *unverified* set. Then, we filtered candidate entities into a manually curated set of *verified* DNA modifications, augmenting them with modification-specific annotations.

The web interface front-end allows users to either search or browse through the catalogue of DNA modifications, integrating ChEBI’s information with our own.

### Identifying candidate DNA modifications from ChEBI

DNAmod leverages ChEBI^21^ to define a set of modified DNA candidates for inclusion and to add preliminary information for each candidate. ChEBI is a database of small biologically relevant molecules, which affect living organisms. We queried ChEBI via ChEBI Web Services ^21^. We used Biopython^9^ and the Python Simple Object Access Protocol (SOAP) client, suds^33^, to query ChEBI and construct the DNAmod database.

ChEBI provides an ontology which encodes the relationships between its compounds. We used this ontology to precisely define the notion of parents and children, which we used to hierarchically retrieve and display modifications. We used two kinds of relationships for this purpose, both of which have associated symbols, defined by ChEBI^12^: 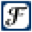 *has functional parent* and Δ*is a*. We used these relationships to find candidate DNA modifications, by identifying entities related to the core nucleobases, which we represent by their symbols: {A, C, G, T, U}. We included uracil, since many of its descendants in the ontology are modifications of thymine (CHEBI:17821, which is equivalent to 5-methyluracil), and are not annotated as descendants of thymine itself. For each of these bases, we imported all entities that are annotated in the ontology as a child of one of these bases, via the 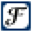 *has functional parent* relationship. ChEBI ranks entities based on their degree of curation. We only imported entities with the highest rating—three stars—indicating manual curation by ChEBI. Whenever possible, we only included entities as nitrogenous bases (nucleobases). If ChEBI did not have the nucleobase, we then selected the nucleoside form and finally, if necessary, the nucleotide. These imported bases formed the candidate set of modifications (the unverified set), from which we created a curated set of DNA modifications (the verified set).

The ChEBI ontology does not generally encode 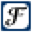 *has functional parent* relationships for nucleobases beyond the children of the unmodified nucleobases. It instead encodes modified nucleobases with an Δ *is a* relationship to their parent base. This is because descendant entities of specific modifications are generally subtypes of the class of modifications from which they originate. For example, 3-methyladenine Δ *is a* methyladenine. Methyladenine, however, 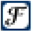 *has functional parent* adenine, since it is conceived of as possessing adenine as a characteristic group and as being derived via functional modification^12^. We therefore need to use both of these relationships, within the ChEBI ontology, to accurately capture the full nucleobase hierarchy.

ChEBI also provides selected citations, associated with some of its entities. We retrieved the citations from ChEBI as PubMed IDs^30^. We used the Biopython^9^ package Bio.Entrez to query the PubMed citation database, using NCBI’s Entrez Programming Utilities^30^. We retrieved the details of each citation, and use them to construct a formatted citation. We currently support only publications indexed in PubMed.

### Manual curation and annotation

We manually created a *whitelist*, which contains our curated (or verified) set of candidates that we deem DNA modifications. For each of these bases, we also imported all descendants with an eventual 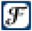 *has functional parent* or Δ *is a* relationship with any of the members of the verified set. We expanded the verified set to include any bases recursively imported in this manner, since they were children of verified DNA nucleobases. This rule had one exception: we excluded any bases that possess an ancestor in our *blacklist* of non–DNA modifications.

We can formalize the above description of bases imported from the ChEBI ontology^12^ and subsequent filtering as follows. Let *a 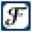 b* specify that *a* has t he 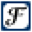 *has functional parent* relationship with *b*. The definition of 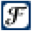 is transitive: for all *n* entities, *l*_*i*_, for *i* = 0 to *n -* 1, between *a* and *b*,

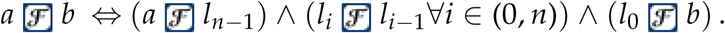

The analogous definitions hold f or Δ.

We call each *l*_*i*_ a *child* of *l*_*i-*1_ and call each *l*_*i-*1_ a *parent* of *l*_*i*_. We refer to *a* as a *descendant* of *b* and refer to *b* as an *ancestor* of *a*. Let 𝒞 represent the first level of children of the unmodified nucleobases, such that 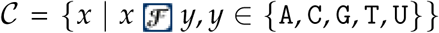. 𝒱 ⊂ 𝒞 Let represent the manually-annotated, verified proper subset of 𝒞.

We manually curated a blacklist of excluded entities, ℬ, satisfying: *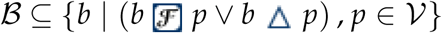.* We imported the set of verified DNA modifications, ℳ, defined in set-builder notation with predicates, as:

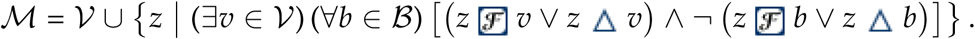

Finally, we added a small number of bases manually, that do not have any of the DNA bases or uracil as a parent in their ontology, but are nonetheless notable modified bases, such as 2^′^-deoxyinosine.

We additionally provided two kinds of manual annotations: sequencing techniques and occurrence in nature, for each modified DNA base. We surveyed the literature of sequencing methods for covalent DNA modifications^5,28,35,37,43^, and annotated the available methods for each base, providing curated citations. These annotations include the method’s name, our categorizations of the basis for the method (such as chemical conversion), its resolution, and any further qualifier (Table 1A). Qualifiers include limitations (such as applicability to only some genomic regions), enrichment methods, and advantages (such as optimization for single-cell sequencing). We considered any method which involves affinity-based recognition of targets to be of “low” resolution^4^. These methods can also suffer from low specificity or antibody cross-reactivity^5^. Conversely, we annotated any methods based principally upon the detection of a chemically converted modification as “high” resolution. This generally reflects the resulting resolution of the method’s output data and often corresponds to the necessity to bin genomic regions during downstream analyses of the detected analyte.

**Table 1.** Possible annotations within DNAmod’s curated (A) sequencing method data and (B) natural occurrence information. Each row lists a field and all terms ever used to annotate it. [square brackets]: optional prefixes. ⟨ angle brackets ⟩: description of term, rather than the complete enumeration provided for other terms.

**(A) Sequencing method annotations**
| Field | Terms |
| --- | --- |
| Mapping method | $\langle$ method abbreviation $\rangle$ |
| Method detail | affinity-based, chemical conversion, chemical conversion and immunoprecipitation, chemical tagging, direct detection, DNMT1 conversion, enzyme-mediated chemical tagging, excision repair enzyme-based, restriction endonuclease |
| Resolution | low, high, single-base |
| Qualifier | 5hmU:G mismatch only, CpG contexts only, [low-input or] single-cell, [methylation-insensitive] restriction digestion, microarray probes, salt gradient stratification, specific fragments, strand-specific, target sequences |

**(B) Natural occurrence annotations**
| Field | Terms |
| --- | --- |
| Function<sup>a</sup> | damage, demethylation intermediate, [possible] epigenetic mark, hypermodified nucleobase, restriction-modification |
| Functional detail | [highly] cytotoxic, mutagenic, reactive oxygen species, specific transcriptional roles, transcription terminator |
| Origin | natural, synthetic, synthetic and RNA |
| Organism | $\langle$ binomial name $\rangle$ |

<sup>a</sup> Each row contains all possible instantiations of the field on the left, except that terms within the “Function” field are often combined, as conjunctions.

For each modified base, we investigated if it had been previously reported to occur *in vivo*. This included any endogenous occurrences, as well as those stimulated exogenously, such as from exposure to an environmental toxin. We annotated any modification observed *in vivo* as “natural”. We additionally provided non-exhaustive examples of some organisms in which the modifications have been reported. We based these annotations on our ability to find evidence of *in vivo* occurrence, as opposed to publications describing only the synthesis or physicochemical properties of a nucleobase. For each of these annotations, we also briefly annotated a primary biological function, if known (Table 1B). For any modification not observed *in vivo*, we annotated it as “synthetic” and listed a reference pertaining to its synthesis or in which the synthetic base was used.

We entered these annotations in two annotation source files (Table 1), which we later imported into our database. This decoupled them from the rest of our pipeline and allows outside experts to submit additions without requiring knowledge of our pipeline or programming workflow.

DNAmod integrates manually-curated nomenclature, including the name and abbreviation deemed most consistent and in common use^8,10,27^. We additionally provide recommendations for one-letter symbols of selected modified bases, and in some instances for their base-pairing complements, as previously described^47^. The DNAmod web interface displays recommended notation in an organized table (Figure 1).

**Figure 1.**
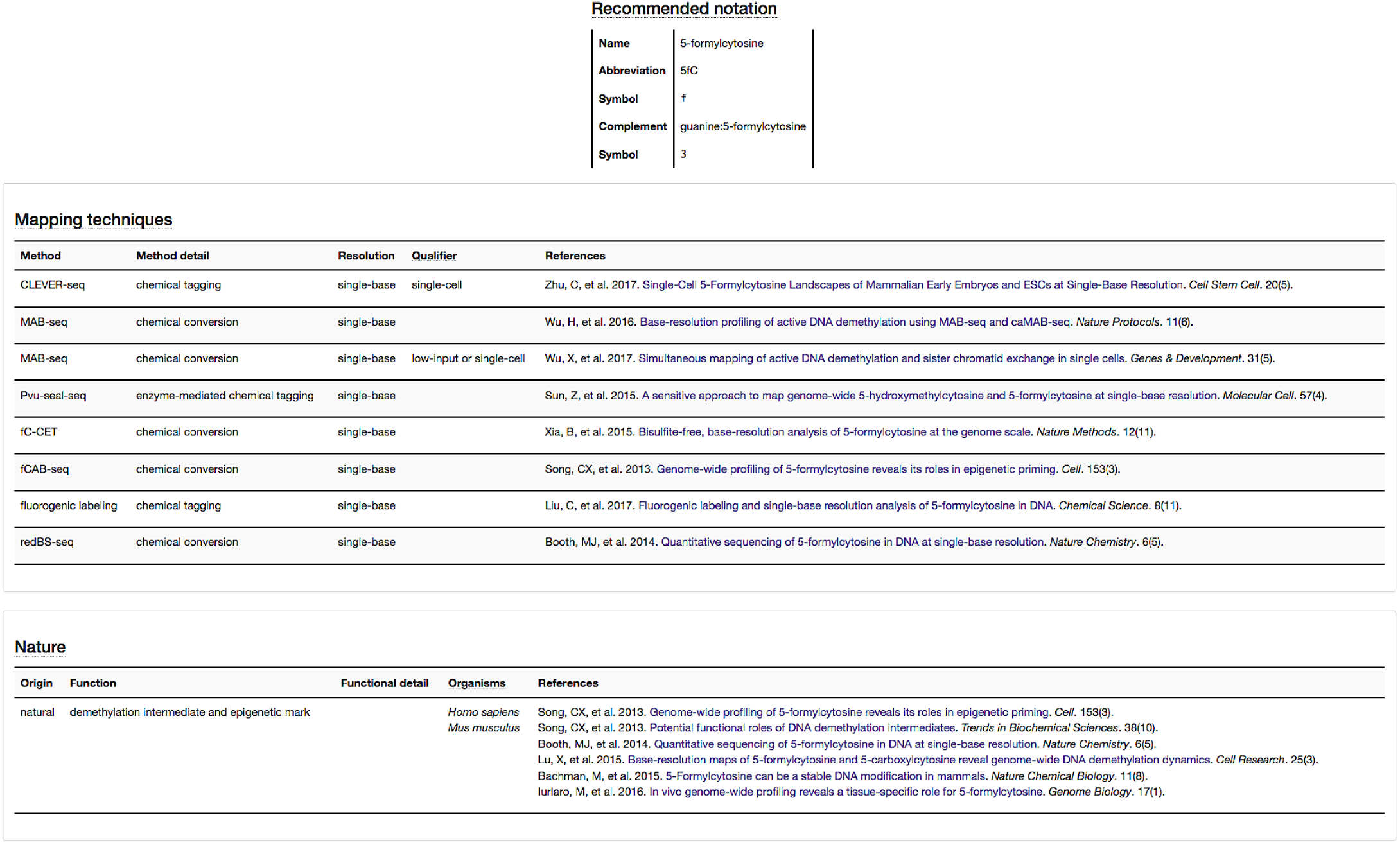
Manually-curated recommended notation, mapping techniques, and natural occurrence data for 5-formylcytosine (5fC). See Table 1 for an explanation of the mapping and natural occurrence table headers.

We store all data, either imported from ChEBI or from our manual annotations, within a SQLite^24^ database, used via the Python sqlite3 package^15^.

### Website generation

We created a static website to display and provide navigation for the information contained within the database. We generated it by formatting the database content using the templating engine Jinja2^40^. Two templates were sufficient to generate all HTML files. We used a single template for all modification pages and another for the homepage. We also recorded the date of the most recent update to the database. All web pages use the Bootstrap^34^ framework, which provides a standardized, portable, and mobile-compatible viewing format. We visualized the chemical structure of each compound from its Simplified Molecular-Input Line-Entry System (SMILES)^50^ data, if available from ChEBI, as a vector graphic. We did this using the cheminformatics toolkit Open Babel^32^, via its Python wrapper Pybel^31^.

### Searching and navigation

DNAmod makes modifications accessible via three main navigation options, each provided on a tab of the DNAmod homepage. First, users may search for modifications by several fields. Second, users may find curated DNA modifications via a pie menu^6^. Third, users may find candidate entities as a list, categorized by their parent unmodified nucleobases.

Client-side search functionality provides a means of rapidly finding bases with differing nomenclature (Figure 2A), while maintaining a static web page. This functionality relies on the elasticlunr.js JavaScript module^45^. Searches match to multiple fields: common or International Union of Pure and Applied Chemistry (IUPAC) names, all synonyms, any assigned abbreviation, and recommended notation symbol, when available. DNAmod displays curated DNA modifications in green, and others in magenta. The search results provide the field matched by the query, such as “abbreviation”, along with the common name of the associated hit.

**Figure 2.**
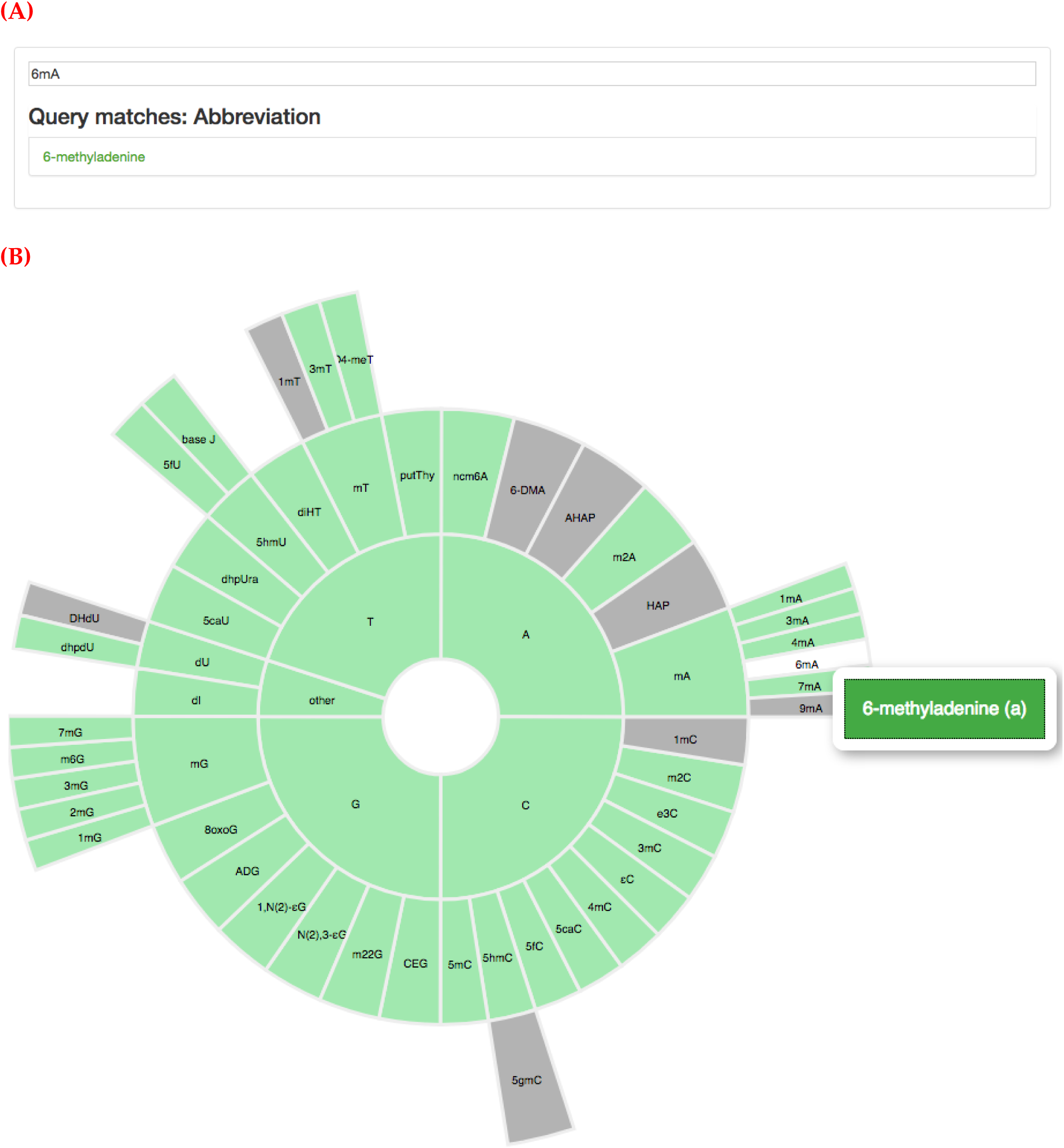
Finding 6-methyladenine by (A) searching for its abbreviation “6mA” or (B) via the pie menu.

Alternatively, users may browse the modifications in DNAmod through a pie menu^6^ interface (Figure 2B). This interface hierarchically arranges the bases according to their structure within the ChEBI ontology. The innermost ring consists of the four unmodified DNA bases, with an additional “other” category. This category encapsulates modified bases found in DNA, but which are not modifications of one of the four DNA bases. Consecutive outer rings represent children of the previous base or category. We demarcated natural versus synthetic bases by colouring natural bases in teal and synthetic bases in grey.

## DNAmod structure and content

Individual modification pages visually represent the data contained within the backing database. We standardize and display all modifications in an identical format. DNAmod may omit some information, however, depending upon the extent of ChEBI’s annotations and whether the page describes a verified DNA modification or merely a candidate entry.

Modification pages begin with a header displaying the DNA modification’s ChEBI name. The top-right corner of the page lists the unmodified ancestor of the modification. For example, 5-hydroxymethyluracil is a modification of thymine (Figure 3), whereas 6-dimethyladenine is a modification of adenine.

**Figure 3.**
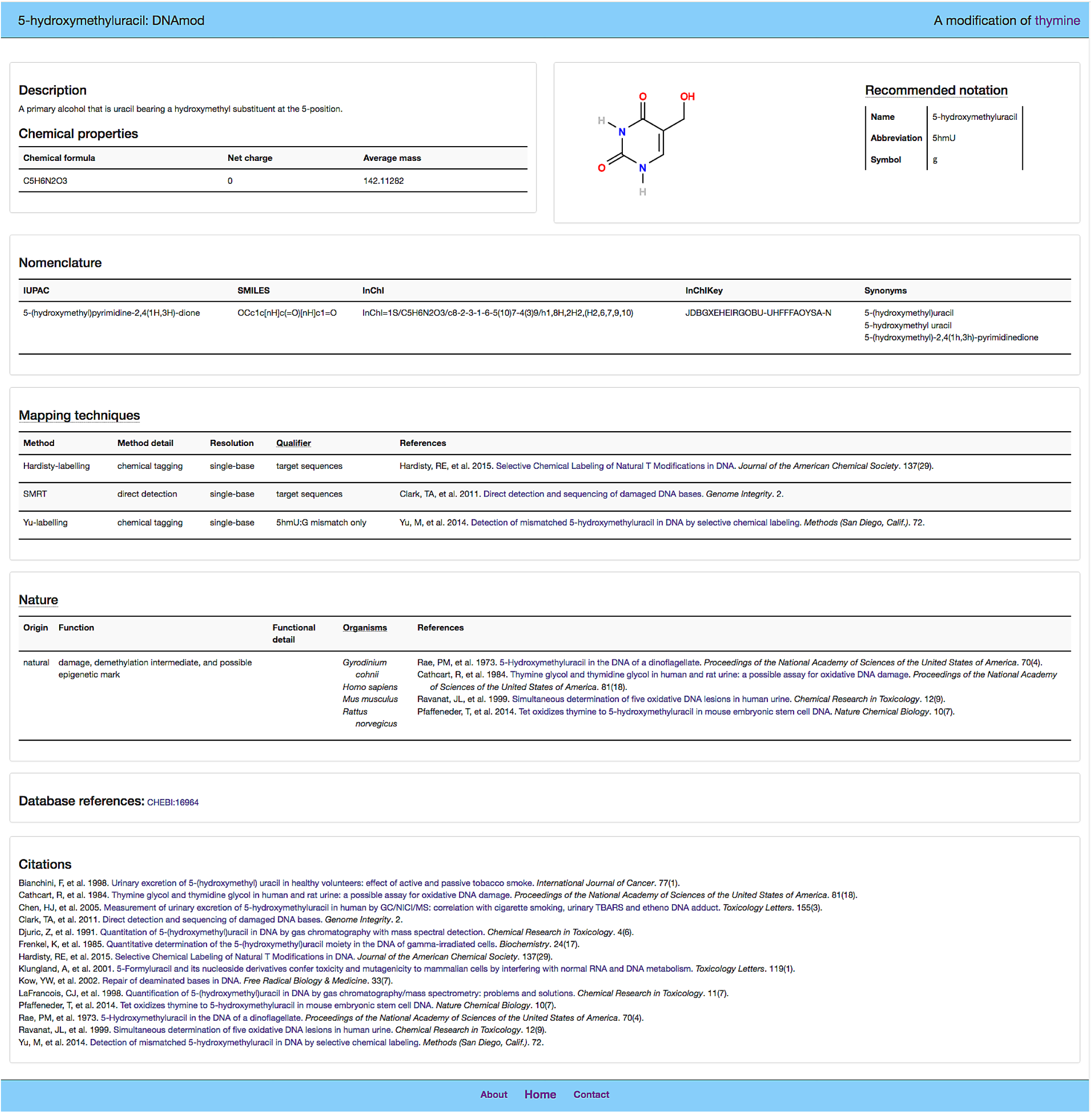
The full modification page for 5-hydroxymethyluracil (5hmU).

Each modification begins with a short textual description of its chemistry, followed by a table containing its chemical properties. We import these from ChEBI, which provides their chemical formula, net charge, and average mass.

We annotate entities with all names available from ChEBI, including: their IUPAC name, SMILES^50^ string, International Chemical Identifier (InChI) and hashed InChIKey^22^ strings, and common synonyms. We also provide a recommended abbreviation and in some instances a suggested single-letter symbol for bioinformatic purposes, from our proposed expanded alphabet^47^ (Figure 3).

We provide literature annotations for many DNA modifications, focusing upon those observed *in vivo*. We provide a list of methods that have been used to map the genomic locations of a modification (see above). We additionally provide information on a modification’s occurrence, either naturally or only synthetically, where applicable, including some organisms in which it has been observed *in vivo* (see above). Finally, each page ends with the ChEBI database reference and a ChEBI-derived list of related literature citations (Figure 3). Our website has semantic web support, making use of the Resource Description Framework in Attributes (RDFa)^38^ technique, augmented by Chemical Information Ontology (CHEMINF)^19^ and PubChemRDF^14^ Semanticscience Integrated Ontology (SIO)^13^ annotations—providing machine-readable descriptions of key website features.

## Discussion

DNAmod enables researchers to rapidly obtain information on covalently modified DNA nucleobases and assist those interested in profiling a modification. It additionally provides a reference toward standardization of modified base nomenclature and offers the potential to track recent developments within the field. We have kept DNAmod up to date for 3 yr and expect to continue to maintain it, particularly as new discoveries about DNA modifications are made. We also hope that DNAmod will serve to highlight underappreciated modifications that may have substantial biological importance.

The nomenclature used to describe a particular DNA modification is often inconsistent, with some early efforts toward standardization of particular classes^10,27^. The ChEBI name, for instance, often corresponds to the common chemical name of the compound, which is occasionally distinct from its common name within the biological literature, in the context of a DNA modification. We address this and attempt to encourage standardization by endeavouring to ensure that other names are annotated, while providing specific nomenclature recommendations. In particular, the suggested name of verified DNA modifications, as displayed on the homepage and within the recommended notation section, is always manually-curated and sometimes differs from the name assigned by ChEBI.

Our database, like many others, relies upon the ChEBI ontology. Like any large and complex endeavour, curating ChEBI is a substantial undertaking, requiring protracted deployment of expertise and effort. While ChEBI has a dedicated team of expert curators, who assiduously and continually improve ChEBI, their resources are naturally limited. Accordingly, while ChEBI has an issue tracker where we and others can suggest changes, revisions to ChEBI are highly dependent on user reports and the team’s available bandwidth. A recent study found that ChEBI contains a non-negligible fraction of errors and omissions, across most entity categories^53^. Such errors naturally propagate to its downstream databases, including our own. While we have made efforts to further curate data and report relevant issues back upstream, we do inherit some errors and limitations. As in any project of this nature, we surely have our own errors and omissions. We lack a dedicated curator; accordingly, we curate this data on a best-effort basis. DNAmod has its own issue tracker, and we would appreciate if users could report any of our own errors or omissions, so that we can address them or facilitate reporting them upstream.

The inclusion of assays available to sequence different DNA modifications provides a means of assessing and selecting a sequencing method. It additionally attempts to track sequencing methods over time, as resolution improves, and especially to highlight recent developments, like direct-detection of various modifications via nanopore sequencing^48^. The sequencing annotations we provide annotate nucleobases which are directly elucidated by the method and only for the base or set of bases which the method independently maps. This includes those that are obtained in addition to another nucleobase. For instance, confounded mixtures are often obtained. For example, 5mC and 5hmC cannot be distinguished with only conventional bisulfite sequencing. Alternatively, some methods have the capacity to independently resolve between modifications, such as various nanopore-based methods. Therefore, while many use oxidative bisulfite sequencing (oxBS-seq) in combination with conventional bisulfite sequencing to elucidate 5hmC via subtraction, we only annotated it as a sequencing method for 5mC, which it directly elucidates^5^. Conversely, we only annotate TET-assisted bisulfite sequencing (TAB-seq) under 5hmC, which it directly elucidates^5^, although many use it to also detect 5mC.

We demarcated bases found to occur *in vivo*, providing examples of organisms in which a modification has been found, along with associated citations. This merely substantiates its *in vivo* presence, however. We did not attempt to comprehensively list the organisms which contain any particular modification. Finally, we expect our brief annotations of the biological roles of various DNA modifications to change as further research is conducted.

### Future work

We plan to keep DNAmod updated continuously, manually reviewing newly added ChEBI compounds, requesting appropriate additions to ChEBI, and curating any improvements. We also endeavour to annotate recently developed sequencing methods as we come across them.

Integrating additional external databases will further increase DNAmod’s utility. In particular, we envision potential integration with domain-specific DNA modification databases, such as those cataloguing compounds formed from the operation of particular biological pathways. For instance, modifications involved in DNA damage and repair could be linked to REPAIRtoire^29^ data.

We used ^ChEBI Web Services 21^ to obtain information from their database. ChEBI has, however, recently released a Python application programming interface (API), permitting us to directly access their data^46^. Switching from our current web-based queries to use of their API would likely result in a more robust system and expedite the database-building process.

## Availability of data and materials

The DNAmod website, including a description and contact information, as well as the backing SQLite database, are freely available at: https://dnamod.hoffmanlab.org. Python source code, web assets, and an issue tracker for this project are available at: https://bitbucket.org/hoffmanlab/dnamod. Persistent availability is ensured by Zenodo, in which we have deposited the current version of our code (https://doi.org/10.5281/zenodo.640631) and SQLite database (https://doi.org/10.5281/zenodo.640561). All source code and web assets are licensed under a GNU General Public License, version 2 (GPLv2). DNAmod’s data is licensed under a Creative Commons Attribution 4.0 International license (CC BY 4.0).

## List of abbreviations

5caC: 5-carboxylcytosine
5fC: 5-formylcytosine
5hmC: 5-hydroxymethylcytosine
5hmU: 5-hydroxymethyluracil
5mC: 5-methylcytosine
6mA: 6-methyladenine
API: application programming interface
ChEBI: Chemical Entities of Biological Interest
CHEMINF: Chemical Information Ontology
DNMT: DNA methyltransferase
InChI: International Chemical Identifier
IUPAC: International Union of Pure and Applied Chemistry
oxBS-seq: oxidative bisulfite sequencing
RDFa: Resource Description Framework in Attributes
SIO: Semanticscience Integrated Ontology
SMILES: Simplified Molecular-Input Line-Entry System
TAB-seq: TET-assisted bisulfite sequencing
TET: Ten-eleven translocation enzyme

## Authors’ contributions

Conceptualization, M.M.H; Methodology, A.J.S., C.V., and M.M.H; Software, A.J.S. and C.V.; Resources, M.M.H; Data Curation, A.J.S. and C.V.; Writing — Original Draft, A.J.S. and C.V.; Writing — Review & Editing, A.J.S., C.V., and M.M.H; Visualization, A.J.S., C.V., and M.M.H; Funding Acquisition, M.M.H; Supervision, C.V. and M.M.H.

### Acknowledgments

We thank Daniel D. De Carvalho and Christopher E. Mason for helpful feedback on early versions of DNAmod. We thank the creators of ChEBI 12, and all those who have worked to improve it 20,21,46. In particular, we thank Gareth Owen, Steve Turner, and Marcus Ennis for actively responding to curation requests and Venkatesh Muthukrishnan for managing ChEBI issues. We thank Egon L. Willighagen for useful suggestions in a PubPeer review of an early version of this work. We thank Carl Virtanen, Qun Jin, and Zhibin Lu for technical assistance.

